# Assessing the role of cold-shock protein C: A novel regulator of *Acinetobacter baumannii* biofilm formation and virulence

**DOI:** 10.1101/2021.05.13.444116

**Authors:** Brooke R. Tomlinson, Grant A. Denham, Robert S. Brzozowski, Jessie L. Allen, Prahathees J. Eswara, Lindsey N. Shaw

**Affiliations:** Department of Cell Biology, Microbiology and Molecular Biology, University of South Florida, 4202 East Fowler Avenue, ISA 2015, Tampa, FL 33620-5150, USA

**Author notes:** **Correspondence:** Lindsey N. Shaw.

## Abstract

*Acinetobacter baumannii* is a formidable opportunistic pathogen that is notoriously difficult to eradicate from hospital settings and can spread quickly via healthcare personnel despite preventive measures. This resilience is often attributed to a proclivity for biofilm formation, which grants *A. baumannii* a higher tolerance towards external stress, desiccation, and antimicrobials. Despite this, little is known regarding the mechanisms orchestrating *A. baumannii* biofilm formation. Herein, we performed RNA-seq on biofilm and planktonic populations for the multidrug resistant isolate, AB5075, and identified 438 genes with altered expression. To assess the potential role of genes upregulated within biofilms, we tested the biofilm forming capacity of their respective mutants from an *A. baumannii* transposon library. In so doing, we uncovered 24 genes whose disruption led to reduced biofilm formation. One such element, cold shock protein C (*cspC*), produced a mucoidal, non-mucoviscous colony phenotype. RNA-sequencing of this mutant revealed the down regulation of pili and fimbriae in the *cspC* mutant, which would explain the decreased biofilm observed. Using MIC assays, we note that the mutant displayed increased antibiotic tolerance that we hypothesize is mediated by overexpression of multi-drug efflux pumps and altered mRNA stability of their corresponding transcriptional repressor. Finally, we show that CspC is required for survival during oxidative stress and challenge by the human immune system, and plays a pivotal role during systemic infection. Collectively, our work identifies a cadre of new biofilm associated genes within *A. baumannii* and provides insight into the global regulatory network of this emerging human pathogen.

## Introduction

*Acinetobacter baumannii* is formidable pathogen that causes over 8,000 cases of multidrug resistant infections annually in the United States and is recognized globally as an urgent health threat (1, 2). This bacterium gained a foothold in the US healthcare system in 2003-2005 when injured service members returned home from the Iraq conflicts harboring *A. baumannii* infections (3), earning it the nickname “Iraqibacter.” Since then, a number of outbreaks have occurred in the US, several of which impacted multiple hospitals (4–6). Fundamental to success of this pathogen is the ability to persist on surfaces for prolonged periods (7–10), the rapid spread via healthcare personnel despite conventional prevention measures (11–13), and its genomic plasticity, driving rapid acquisition of antibiotic resistance (14). Thus, it has been suggested that *A. baumannii* employs a ‘persist and resist’ strategy, as opposed to a particular set of toxins or molecular determinants that may otherwise dictate an organism’s potential for virulence (reviewed in (15)).

Like many bacteria, the resilience of *A. baumannii* is often attributed to a proclivity towards biofilm formation and its ability to respond to external stress (16). Biofilms are multicellular aggregates surrounded by exopolymeric substance (EPS), which consists of extracellular DNA, proteins, and polysaccharides (17–19). Biofilms exhibit greater resistance to antimicrobials due to reduced diffusion of antibiotics through the multilayered matrix (20) and physiological heterogeneity of the bacterial community (21), which ultimately promote infection chronicity. These factors also contribute to tolerance of environmental stress, such as desiccation (16), which significantly enhances *A. baumannii* persistence in hospital settings. Despite this, only a handful of contributing factors and conditions for *A. baumannii* biofilm formation have been identified (reviewed in (22) and (23)). In particular, pili production plays a pivotal role in the first step of biofilm initiation: the irreversible attachment of bacteria to a surface. Two highly conserved, essential *A. baumannii* attachments factors have been identified to date, chaperone-usher pili (Csu) and outer membrane protein A (OmpA) (22, 24, 25). With regards to the latter, OmpA has been shown to facilitate attachment to abiotic surfaces as well as epithelial cells (25). Comparatively, the Csu assembly system (CsuA/B, CsuA-E) seems to have a more critical role mediating attachment to abiotic surfaces and purportedly does so by hydrophobic interaction with structurally variable substrates, including plastic and glass (24, 26). Transcriptional regulation of the *csu* operon is governed by the BfmRS two-component regulatory system (27), although whether this is by direct or indirect means remains unresolved. During sub-MIC antibiotic treatment, BfmRS also enhances the expression of the major capsular biosynthesis gene cluster (K locus) to promote capsule production, another major biofilm determinant (28). Previous work on *Acinetobacter baumannii* and its non-pathogenic relative, *Acinetobacter baylyi,* has shown that capsule production augments desiccation tolerance, which enhances biofilm survival on surfaces for prolonged periods (16, 29, 30). Overall, capsule production is important for *Acinetobacter* desiccation tolerance, biofilm integrity, and persistence, and the regulatory network for capsule production is intertwined with that of pili production and biofilm formation.

To identify novel factors which drive *A. baumannii* biofilm formation, herein we implemented a transcriptomic approach, comparing the global expression profiles of biofilm to that of planktonically growing counterparts. We then identified genes for which transcription was enriched within biofilm, obtained mutants harboring transposons disrupting these genes, and screened for changes in biofilm integrity. In so doing, we identified 24 genes whose disruption resulted in a notable decrease in biofilm mass compared to wildtype. During this screen, we identified one such factor, cold-shock protein C (CspC), disruption of which resulted in a mucoidal, but non-mucoviscous, colony phenotype. Cold-shock proteins (Csps) have a demonstrated importance in cell aggregation (31), extracellular polysaccharide production, and membrane fluidity in other organisms (32). Furthermore, Csps are multifunctional proteins, which can have pleiotropic functions as transcriptional regulators and RNA chaperones (33). Thus, these proteins can have impacts at the transcriptional, post-transcriptional, and translational levels (reviewed in (34, 35)). Although Csps often play important roles in bacterial stress response, they have not been previously implicated in the regulation of *A. baumannii* biofilm formation. In this work, we reveal that disruption of *cspC* leads to enhanced tolerance to polysaccharide degradation, altered antibiotic resistance profiles, and diminished abiotic surface adherence. The *cspC* mutant also exhibited diminished survival in a murine model of infection. Collectively, these findings establish CspC as a novel regulator of *A. baumannii* biofilm formation and disease causation.

## Methods

### Bacterial strains and growth conditions

Bacterial strains used for this study are listed in Table 1*. A. baumannii* and *Escherichia coli* strains were routinely cultured in lysogeny broth (LB) with shaking or on LB agar at 37°C. When appropriate, media was supplemented with tetracycline and hygromycin at a final concentration of 5 μg/mL and 160 μg/mL respectively. *A. baumannii* biofilm formation was assessed as described previously for microtiter plate biofilms (38) with slight modifications. Briefly, overnight cultures were normalized to an OD_600_ of 5.0 in phosphate-buffered saline (PBS), before 20 μL was added to 180 μL of fresh LB in 96-well microtiter plates for a final OD_600_ of 0.5. For assays incorporating strains harboring pMQ557, LB was supplemented with hygromycin.

**Table 1.** Bacterial strains and plasmids used in this study.

| Strains | Description | Source |
| --- | --- | --- |
| ***E. coli*** |  |  |
| DH5α | Cloning strain | (36) |
| ***A. baumannii*** |  |  |
| AB5075 | Parent strain | (37) |
| *cspC::tn* | AB5075 with transposon insertion in ABUW_1377 (*cspC*) | (37) |
| WT | AB5075 containing pMQ557 | This study |
| *cspC*<sup>-</sup> | *cspC::tn* containing pMQ557 | This study |
| *cspC*<sup>+</sup> | *cspC::tn* containing pMQ557::*cspC* | This study |
| **Plasmids** |  |  |
| pMQ557 | Cloning vector for complementation | Gift, Dr. R. Shanks, University of Pittsburgh |
| pLSBT1 | pMQ557::*cspC* | This study |

### Mutant strain confirmation and complement strain construction

Transposon mutant strains were obtained from the *A. baumannii* AB5075 transposon mutant library (37). Transposon insertion was confirmed for the *cspC* transposon mutant by PCR and sequencing using primers OL5087 and OL5088 which flank *cspC* (all primers are listed in Table S1). A *cspC*-complement PCR fragment was amplified using primers OL5088 and OL5214 and cloned into plasmid pMQ557. In addition to the full length of *cspC*, the complement fragment includes 382 nucleotides immediately upstream and 97 nucleotides directly downstream of the *cspC* coding region to ensure the native promoter and full-length mRNA (as revealed by RNA-seq) were incorporated. Complementation plasmid pMQ557::*cspC* was transformed into chemically competent *E. coli* (DH5α) and confirmed using pMQ557 screening primers OL4163 and OL4164 followed by sequencing. Plasmid containing the correct sequence was transformed into the *cspC* transposon mutant strain via electroporation (44) to generate the complement strain, *cspC^+^*. Given that vector is present in *cspC^+^*, the empty pMQ557 vector was also transformed into the wildtype parental strain and *cspC* transposon mutant strain (*cspC*^−^). For all assays *cspC^+^,* wildtype and *cspC* mutant strains bearing pMQ557 were used for comparison throughout.

### RNA sequencing

*A. baumannii* biofilm formation was initiated in 96-well microtiter plates as described above in biological triplicate for 24 h at 37°C in a static incubator. To collect planktonic samples, 75 μL of supernatant was removed from each well and pooled. Planktonic cells were immediately combined with 5 mL of ice-cold PBS, and pelleted by refrigerated centrifugation. For biofilm samples, the remaining supernatant was removed and biofilm containing wells were washed three times with 200 μL of ice-cold PBS. Ice-cold PBS was added a final time and pipetted vigorously to disrupt biofilm cells. Biofilm cells from different wells were pooled, immediately combined with an additional 5 mL of ice-cold PBS, and pelleted by refrigerated centrifugation.

For collection of AB5075 wildtype and AB5075 *cspC::tn* mutant samples, strains were grown in LB overnight with shaking at 37°C in biological triplicate. Overnight cultures were diluted 1:100 into 5 mL of fresh LB, grown to exponential phase, and subsequently used to seed new 100 mL cultures at OD_600_ 0.05. Samples were harvested after 3 hours of growth, added to an equal volume of ice-cold PBS, and pelleted by centrifugation at 4°C.

Total RNA was isolated from cell pellets as described previously (45) using an RNeasy Kit (Qiagen) and DNA removal was performed using a TURBO DNA-free kit (Ambion). DNA removal was confirmed by PCR using OL398 and OL399. Sample quality was assessed using an Agilent 2100 Bioanalyzer system and corresponding RNA 6000 Nano kit (Agilent) to confirm RNA integrity. Samples with a RIN of ≥9.9 were used in this study. Prior to mRNA enrichment, biological triplicates were pooled at equal RNA concentrations. rRNA was then removed using a Ribo-Zero Kit for Gram Negative Bacteria (Illumina) followed by a MICROBExpress Bacterial mRNA enrichment kit (Agilent). Removal efficiency of rRNA was confirmed using an Agilent 2100 Bioanalyzer system and RNA 6000 Nano kit (Agilent). Enriched mRNA samples were then used for RNA sequencing using an Illumina NextSeq. Library preparation and RNA sequencing was performed following Truseq Stranded mRNA Kit (Illumina) recommendations omitting mRNA enrichment steps. Quality, concentration, and average fragment size of each sample was assessed using an Agilent 2100 Bioanalyzer system and RNA 6000 Nano kit (Agilent) prior to sequencing. Library concentration for pooling of barcoded samples was assessed by RT-qPCR with a KAPA Library Quantification kit (KAPA Biosystems) as recommended for high sensitivity. Samples were run on an Illumina NextSeq with a corresponding 150-cycle NextSeq Mid Output Kit v2.5. Experimental data from this study were deposited in the NCBI Gene Expression Omnibus (GEO) database (GEO accession numbers GSE164233 and GSE164290).

### RNAseq bioinformatics

Data was exported from BaseSpace (Illumina) in fastq format and analyzed using CLC Genomics Workbench 20 (Qiagen Bioinformatics). Reads were imported and failed reads were removed using the Illumina Paired Importer tool, with quality score parameter option set to Illumina Pipelines 1.8 and later. The total number of reads generated for each sample was at least 15.49 million and up to 22.22 million, resulting in ≥545x read coverage for each sample. Reads corresponding to rRNA were filtered, removed by aligning to known rRNA sequences, and discarded. Samples contained between 0.27% and 0.71% rRNA. Remaining read sequences were aligned using the RNA-seq Analysis tool (v0.1) with default parameters and defined strand specificity to the *A. baumannii* AB5075 NCBI reference genome (CP008706.1). Gene expression values were calculated using the Expression Browser tool (v1.1) specifying transcripts per million (TPM) as the output. Differential expression values between samples were generated using the Differential Expression in Two Groups tool (v1.1) for whole transcriptome RNA-seq samples. Differential expression is reported as fold-change of expression for biofilm relative to planktonic samples, and cspC mutant relative to wildtype samples. Library size normalization is automatically performed using the trimmed mean of M values (TMM) method by the Differential Expression in Two Groups tool (46). Ontology classification of genes was assigned based on the Kyoto Encyclopedia of Genes and Genome (KEGG) (47). Genomes and differential expression visualizations (Fig. 1A and 5A) were generated using Circos (48).

**Figure 1.**
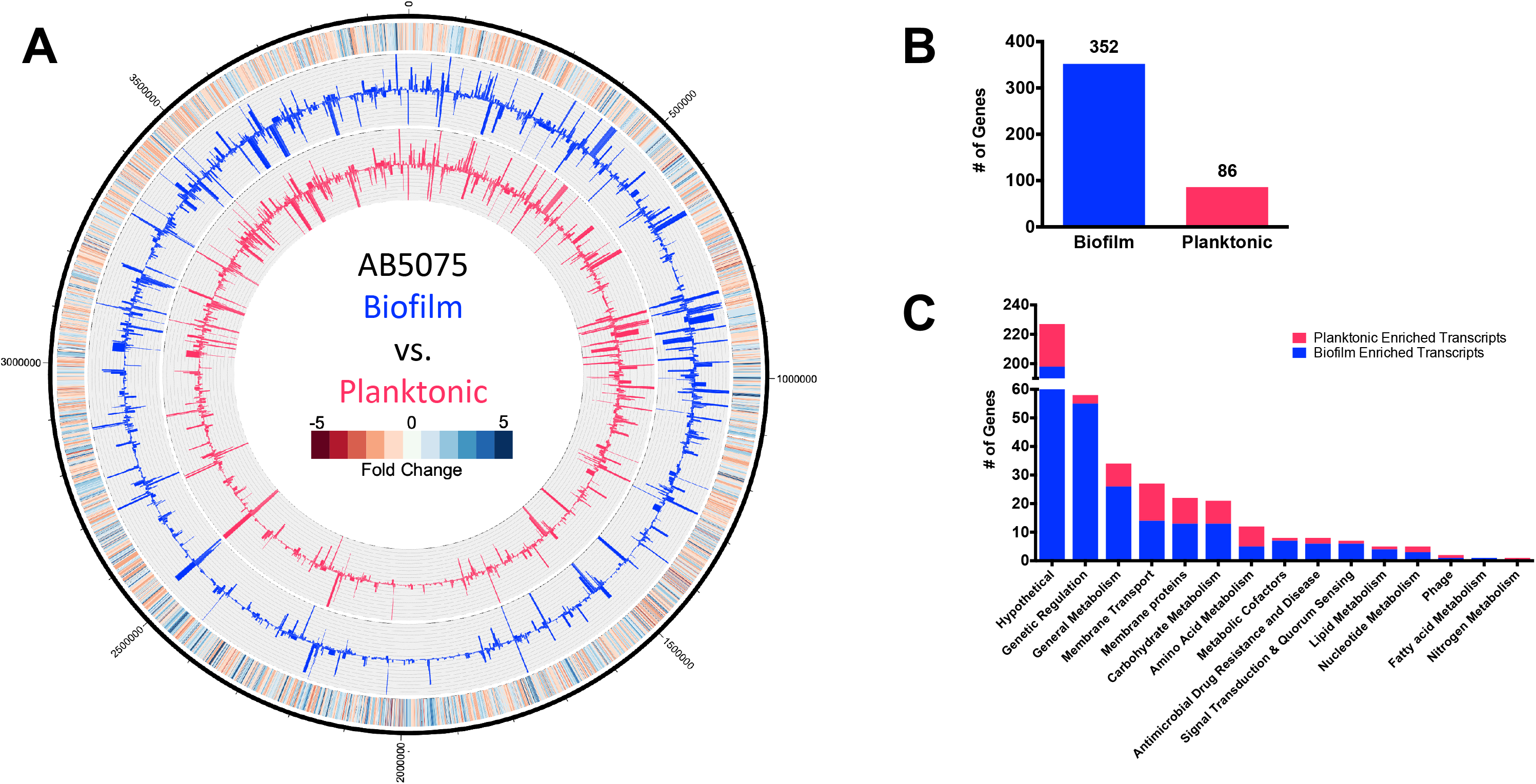
*A. baumannii* biofilms exhibit differential expression patterns compared to planktonic cell populations. The genomic map (**A**) depicts changes in planktonic (inner circle, pink) and biofilm (outer circle, blue) transcriptomes, reported as TPM expression values. The outermost circle is a heat map demonstrating fold change in expression, where red or blue indicates higher expression in the biofilm or planktonic cells, respectively. The number of genes that were preferentially expressed ≥2-fold in biofilm (blue) or planktonic (pink) cell populations are tallied (**B**) and further sorted by function based on KEGG ontology (**C**).

### RT-qPCR transcriptional analysis

To validate RNA-seq findings, a selection of genes were assayed by Real-Time Quantitative Reverse Transcription PCR (RT-qPCR). Strains were grown and RNA harvested as described above for RNA-seq studies performed on biofilm and planktonic samples in biological triplicate. Total RNA was isolated from cell pellets and DNA removal was performed as described above. Sample quality was assessed using an Agilent 2100 Bioanalyzer system and corresponding RNA 6000 Nano kit (Agilent) to determine RNA concentration and to confirm RNA integrity. Samples with a RIN of ≥9.9 were used. One microgram from each sample was reverse transcribed using an iScript cDNA Synthesis Kit (BioRad). RT-qPCR was then performed using gene-specific primers (Table S1) and TB Green Premix Ex Taq (Takara). Levels of gene expression were normalized to that of 16S ribosomal RNA gene (OL4498 and OL4449) and fold-change of expression was assessed for biofilm relative to planktonic samples, and cspC mutant relative to wildtype samples, using the 2^−ΔΔCt^ method (49).

Similarly, transcriptional analysis was used to assess changes in *cspC* transcript levels under cold stress. *A. baumannii* wildtype cultures were grown overnight as described above, in biological triplicate, and sub-cultured into 5 mL of fresh media. After 3 hours of growth at 37°C, new 5 mL cultures were seeded and standardized to and OD_600_ of 0.05. Bacteria were transitioned to 15°C for 15 minutes to induce cold shock, and control samples were left at 37°C. Following this, samples were combined with an equal volume of ice-cold PBS and pelleted by centrifugation at 4°C. Total RNA isolation, DNA removal, and reverse transcription were performed as described above. Quantitative, real-time RT-PCR (qRT-PCR) was then performed using *cspC*-specific primers OL5216 and OL5217, and TB Green Premix Ex Taq (Takara). Levels of gene expression were normalized to that of 16S ribosomal RNA gene (OL4498 and OL4449) and fold-change of expression was assessed for cold shocked samples relative to non-cold shocked samples using the 2^−ΔΔCt^ method (49).

### Crystal violet and real-time biofilm assays

*A. baumannii* biofilm formation was performed as described above in biological triplicate. After 24 hours of static growth, biofilms were washed 3 times with PBS and fixed with 100 μL of 100% ethanol. After drying, 200 μL of crystal violet was added, incubated at room temperature for 15 minutes, and biofilms washed 3 times with PBS. After a second drying step, 100 μL of 100% ethanol was added to solubilize the crystal violet. Absorbance of solubilized crystal violet was measured at OD_543_ and reported as percent variance to that of the wildtype strain.

A Real Time Cell Analyzer (RCTA) xCELLigence MP (ADCEA Bioscience) instrument was used to monitor biofilm formation over time. The xCELLigence RTCA MP was placed in a 37°C incubator for one hour prior to experimentation to allow the instrument temperature to equilibrate. Next, 96-well E-plates were loaded with 180 μL of LB, positioned in the RTCA, and measured for background signal. Using the same plate, *A. baumannii* biofilms were prepared as described above, and statically incubated in the RTCA, with reads taken every 15 minutes for 25 h. The data generated herein is from nine biological replicates per strain.

### Microscopy

Cell morphology was assessed by fluorescence microscopy as previously described (50) with minor modifications. Briefly, single 24h wildtype, *cspC::tn* mutant and complemented strain colonies were resuspended in 100 µL of 1x PBS. Cell membranes and DNA was stained with FM4-64 and DAPI, respectively, at final concentrations of 1 µg/mL. Cell suspension (5 µL) were spotted onto a glass coverslip of glass bottom dishes (MatTek) and subsequently covered with a sterile pad of 1% agarose in water. Imaging was completed at room temperature inside a DeltaVision Elite deconvolution fluorescence microscope (GE Applied Precision) environmental chamber. Photos were captured using a CoolSNAP HQ2 camera (Photometrics) and images were acquired by taking 17 z-stacks at 200 nm intervals. All images were deconvolved using the softWoRx (GE Applied Precision) imaging software.

### Extracellular DNA assays

Extracellular DNA production of *A. baumannii* biofilms was analyzed quantitatively as described previously (51). Briefly, *A. baumannii* biofilm formation was initiated as described above in biological triplicate. After 24 hours of growth, supernatant was removed, and biofilms were washed once with 200 μL PBS. eDNA in biofilms was quantified by Quant-iT PicoGreen dsDNA labelling (Thermo Fisher) and fluorescence measured using a Synergy2 plate reader (BioTek).

### Biofilm inhibition by proteinase K and sodium *meta*-periodate

Disruption of *A. baumannii* biofilms by sodium *meta-*periodate and proteinase K was analyzed quantitatively as described previously (52). Briefly, *A. baumannii* biofilm formation was initiated as described above in biological triplicate. Biofilms were supplemented with a final concentration of 100, 50, 25, 12.5, 6.25, 3.13, 1.56, 0.78, or 0 mM of sodium *meta*-periodate. Alternatively, biofilms were supplemented with a final concentration of 50 μg/mL of proteinase K. Biofilms were allowed to form for 24 h at 37°C and then quantified by crystal violet assay as described above.

### Cold-shock recovery and survival

*A. baumannii* cultures were grown overnight, in biological triplicate, and sub-cultured into 5 mL of fresh media. After 3 hours of growth, new cultures were seeded and standardized to and OD_600_ of 0.05. For cold stress recovery, after one hour of growth at 37°C bacteria were transitioned to 15°C for 1 hour. Following this cold stress, bacteria were returned to 37°C with shaking. Bacteria were plated on TSA every hour to determine CFU/mL as a measure of recovery rate. Alternatively, for cold stress survival, bacteria were immediately placed at 15°C after culture standardization and plated on TSA every hour to determine CFU/mL for monitoring survival.

### Antibiotic susceptibility assays

Antibiotic sensitivity was assessed by performing minimum inhibitory concentration (MIC) assays as previously described (53). Briefly, *A. baumannii* strains were grown in LB overnight, in biological triplicate, at 37°C with shaking. Overnight cultures were diluted 1:1,000 with fresh LB and 195 μL was added to 96-well microtiter plates. Subsequently, antibiotics were serial diluted and 5 μL of each concentration, or solvent (no-treatment control), was added. Antibiotic solvents were as follows: ciprofloxacin, 0.1M NaOH; chloramphenicol, 70% EtOH; streptomycin, H_2_O; gentamicin, 100% EtOH; kanamycin, H_2_O; neomycin, H_2_O; Fosfomycin, H_2_O; oxacillin, H_2_O. Cultures were grown overnight at 37°C with shaking. MIC is reported as the lowest antibiotic concentration resulting in inhibition of growth compared to no-treatment control.

### CspC protein architecture analysis

Domain and motif scanning was performed using ScanProsite (54) with the amino acid protein sequence of CspC (GenBank: AKA31122.1) as the input. The alignment of 207 UniProtKB/Swiss-Prot sequences of true positive hits for the detected cold shock domain profile (PS51857) were retrieved and a sequence logo was generated from this alignment on Prosite. Three-dimensional protein modeling via homology modelling was completed using Swiss-Model (55). The Swiss-Model template library contained 244 templates matching the CspC amino acid sequence, with the most closely related, and the sequence with the highest global model quality estimate, being CspA of *Escherichia coli* (56).

### Transcriptional arrest and determination of mRNA half-life

Determination of RNA half-life was performed as described previously with minor modifications (57). Six biological replicates of *A. baumannii* cultures were grown overnight and sub-cultured into 5 mL of fresh media. After 3 hours of growth, new cultures were seeded and standardized to and OD_600_ of 0.05 in 100 mL of fresh LB. After 3 h of growth to reach exponential phase, and prior to transcriptional arrest, 5 mL of each culture was collected, immediately combined with 5 mL of ice-cold PBS, and pelleted by refrigerated centrifugation (t=0). Rifampin at a final concentration of 250 μg/mL was then added to bacterial cultures. At 5, 10, 15, 30, and 45 minutes post-treatment, 5 mL of each sample was collected, immediately combined with 5 mL of ice-cold PBS, and pelleted by refrigerated centrifugation. Immediately following each refrigerated centrifugation step, supernatant was removed and cell pellets were stored at −80°C. Total RNA extraction, confirmation of RNA quality, and RT-qPCR was performed as described above for each sample (primers used are listed in Table S1). RNA abundance at each timepoint post-treatment was calculated for each biological replicate and measured in technical triplicate for each timepoint using 2^−ΔΔCT^ relative to initial RNA abundance (t=0). These values were plotted as a function of time and an exponential, one phase decay curve was fitted using GraphPad Prism. The decay curve is represented as R(*t*) = R_0_*e^−kt^*, where R_0_ and R(*t*) are relative RNA abundance at initial and subsequent timepoints, respectively (58). The decay rate constant, *k*, is equal to ln(2)/*t*_1/2_, where *t*_1/2_ is mRNA half-life. Accordingly, half-life was derived from this equation as *t*_1/2_ = ln(2)/*k*.

### RNA secondary structure predictions

RNA secondary structure predictions were generated using RNAfold (ViennaRNA package v2.4.18) using default parameters (59). Full length mRNA sequences, as revealed by RNA-seq read mapping, were used as input. All structures predicted by RNAfold were inspected and compared. Consensus structures with the lowest minimum free energy were downloaded in Vienna format and used to draw RNA structure using the forna software (ViennaRNA package v2.4.18) (60).

### Human blood survival assay

Survival in whole human blood was performed as previously described (61) with minor modifications. Briefly, *A. baumannii* cultures were grown overnight, in biological triplicate, and sub-cultured into 5 mL of fresh LB. After 3 hours of growth, 10 mL of cells were centrifuged, washed with PBS, and diluted to an OD_600_ of 0.5. Cells were then added to 1 mL of deidentified whole human blood (BioIVT) at a final OD_600_ of 0.05. The initial inoculum of each strain was determined at this time by serial dilution and plating on LB agar. Blood cultures were incubated at 37 °C with agitation, and CFU/mL of each strain was determined, every hour for six hours, by serial dilution and plating on LB agar.

### Oxidative stress assay

Oxidative stress was assessed using hydrogen peroxide as described previously (62) with minor modifications. Briefly, *A. baumannii* cultures were grown overnight in biological triplicate and sub-cultured into 5 mL of fresh media. After 3 hours of growth, new cultures were seeded and standardized to and OD_600_ of 0.05 in fresh LB. Hydrogen peroxide was then added to the cell suspensions for a final concentration of 2 mM and grown at 37°C with agitation. Cells were collected (500 µL) at each indicated timepoint and supplemented with catalase (10 µg/mL) to neutralize the effects of hydrogen peroxide. Cells were then serial diluted and plated on LB agar to determine surviving CFU/mL.

### Mouse infection model

The experiments were performed with the prior approval of the University of South Florida Institutional Animal Care and Use Committee. A murine model of dissemination was performed based on previous studies (63, 64). Briefly, 13-week-old, female BL-6 mice were purchased from Charles River Laboratories and allowed to acclimate for 2 weeks prior to the start of experimentation. AB5075 wild-type and *csp::tn* strains were grown in LB as described above in biological triplicate. Overnight cultures were then sub-cultured 1:100 in fresh LB and grown for an additional 3 hours. Cultures were then pelleted by centrifugation, washed twice in PBS, and standardized to an OD_600_ of 5.0. From this resuspension, the average number of CFU/mL was calculated for each strain by plating on LB agar. This was repeated on three separate days using two biological replicates per strain, and the average number of CFU/mL across replicates was calculated for each strain. This average number of CFU/mL was used to determine the volume of bacteria needed to obtain a 5-mL inoculum of 2.5 × 10^8^ ([2.5 × 10^8^/average CFU/mL] × 5 mL). On the day of infection, appropriate aliquots from fresh overnight cultures were prepared in the same manner, diluted to 2.5 × 10^8^ CFU/mL, and 100 μL of suspension was administered to 10 mice per strain via retroorbital injection, providing a final inoculum of 2.5 × 10^7^ CFU/mL. Infections were monitored and the mice were sacrificed 6 hours post infection. At this time, whole liver, heart, kidneys, spleen, lungs, and brain were harvested and immediately stored at −80°C. Organs were individually homogenized using a Bullet Blender (Next Advance) in 1.5 mL PBS and serially diluted onto LB agar to determine bacterial burden (CFU/mL). A Mann-Whitney nonparametric test was performed to determine statistical significance of bacterial burden for each organ between the mutant and wild-type strains.

## Results

### Identification of *A. baumannii* biofilm-associated genes

To inform on factors potentially important to biofilm formation, global gene expression within AB5075 wild type biofilm populations was compared to planktonic counterparts (Fig. 1A, Table S2). Of the genes differentially expressed by a magnitude ≥2-fold, 352 genes were expressed higher in biofilm cell populations (Fig. 1B), whilst just 86 genes were enriched ≥2-fold within planktonic cells.

When sorted ontologically, most biofilm-associated transcripts were categorized within the genetic regulation group; fulfilling roles pertaining to DNA replication, transcriptional regulation, and translation (Fig. 1C). Of the 58 genes within this cluster, 27 are annotated as transcription factors. Furthermore, an overwhelming 25 of these transcription factors were enriched in the biofilm population. The *A. baumannii* strain AB5075 genome encodes for 243 transcription factors (65); indicating 10% of known transcription factors are activated in biofilms, with 1% deactivated, suggesting a distinct global regulatory response within this population. Of the 25 biofilm-associated transcription factors, the majority (n=23) are conserved among *A. baumannii* strains, with one, ferric uptake regulator *fur*, having been previously deemed essential for growth in rich medium (37, 66). Of the two less-conserved regulators, HTH_3 family transcriptional regulator, *ABUW_RS16125,* was previously shown to be essential for survival during *Galleria mellonella* infection (67). This may suggest a dual virulence and biofilm enhancing role for ABUW_RS16125 in *A. baumannii* strains encoding this gene.

The upregulation of a cadre of genes involved in metabolic pathways was also apparent within biofilms. The most drastically enriched transcripts were those responsible for energy metabolism, and specifically the sulfur metabolism pathway (68). This included genes encoding for subunits of the taurine transporter permease (*tauA, tauB,* and *tauC*), taurine dioxygenase (*tauD*), sulfate transporter permeases (*cysT, cysW*), and thiosulfate-binding protein (*ABUW_RS05010*), each of which showed ≥7.69-fold enrichment. Several additional genes involved in the sulfur metabolism pathway were also upregulated by ≥2-fold in biofilms, including sulfate transporter permease (*ABUW_RS04990*), thiosulfate-binding protein (*ABUW_RS01375*), and sulfate adenylyl transferase subunits (*cysN* and *cysD*).

Six genes identified as contributors to antimicrobial drug resistance and disease were also upregulated in biofilm. This group included *ABUW_RS09485* and *ABUW_RS09480*, which encode a putative efflux pump and its accompanying membrane fusion protein component, respectively, and are homologous to the *E. coli* efflux pump, EmrAB. Other ontologies with altered expression in biofilms included additional membrane transporters (e.g. – *ABUW_RS11790, ABUW_RS11795, ABUW_RS18795*) and putative membrane proteins (e.g. – *ABUW_RS10540, ABUW_RS17095, ABUW_RS18900*); carbohydrate metabolism (e.g. – *ABUW_RS08155, leuA, ABUW_RS10665*); and amino acid metabolism (e.g. – *yncA, ABUW_RS13045, mmsB*). Thus, it is clear that the biofilm has unique regulatory networks activated compared to that of planktonic cells, which reflect distinct metabolic needs and response to the environment.

To validate RNA-seq findings, a selection of genes were assayed by Real-Time Quantitative Reverse Transcription PCR (RT-qPCR), with fold-change in expression proving comparable to RNA-seq findings (Fig. S1).

### Physiological impact of biofilm-associated genes on biofilm integrity

Preferential expression within biofilm populations does not necessary indicate that a given gene product has a role in biofilm formation. Thus, to explore genes with a tangible role in biofilm formation, we first narrowed the list of 352 genes with biofilm-enriched transcripts to those meeting the following criteria: confident read coverage (>85% uniquely mapped reads), relatively strong expression (TPM ≥115 in biofilm), and greatest fold-increase (>3-fold) (Table 2). Of the genes fitting this criteria, the most highly expressed gene was *ssrS* (62175.62 TPM), which produces the well-conserved, 6S RNA (69); whilst the most upregulated gene within biofilms compared to planktonic populations was *ABUW_RS05005* (+60.17-fold), which encodes an uncharacterized alpha/beta hydrolase fold protein. We next assessed whether the 40 genes within these parameters have an impact on biofilm formation by performing classic crystal violet biofilm assays on transposon mutants for the genes of interest contained within the University of Washington *A. baumannii* AB5075 transposon mutant library (37). In so doing, we uncovered 24 genes whose disruption led to reduced biofilm formation (Fig. 2). Of these, 17 mutant strains showed >50% reduction in biofilm. The largest changes were observed for *ABUW_RS09460* and *ABUW_RS05005,* with >72% reduction in biofilm mass for their respective mutant strains. Although both *ABUW_RS09460* and *ABUW_RS05005* mutants show comparable reduction in biofilm mass, *ABUW_RS09460* was only modestly upregulated (3.07-fold), whereas *ABUW_RS05005* was markedly upregulated (60.17-fold) in biofilm. Significant changes in biofilm formation were also observed for *cspC*, and *hscB* mutants (>40% reduction), with *cspC* being the highest expressed gene (1324.94 TPM), whilst *hscB* was one of the lowest expressed genes (167.95 TPM) within biofilms investigated in these assays. Altogether, neither fold-change in expression nor magnitude of expression exclusively correlate with physiological impact on biofilm formation, but rather each feature must be considered holistically.

**Table 2.** Prioritized genes exhibiting preferential expression within biofilms.

| Gene | Expression (TPM)<sup>†</sup> | Fold Change<sup>‡</sup> |
| --- | --- | --- |
| *ssrS* | 62175.62 | 3.15 |
| *ffs* | 4153.04 | 3.38 |
| *leuA* | 1513.08 | 3.55 |
| *cspC* (ABUW_RS06720) | 1324.94 | 3.16 |
| *hslR* (ABUW_RS08520) | 1059.73 | 3.24 |
| *ABUW_RS08155* | 1037.86 | 3.58 |
| *ABUW_RS02640* | 737.36 | 3.18 |
| *ABUW_RS09455* | 717.76 | 3.53 |
| *cysD* | 565.84 | 3.95 |
| *ABUW_RS01835* | 554.93 | 3.92 |
| *bfr_2* | 532.31 | 3.36 |
| *ABUW_RS08160* | 511.43 | 4.02 |
| *cysP* (ABUW_RS05010) | 440.27 | 51.8 |
| *iscA* | 370.19 | 3.93 |
| *ABUW_RS20285* | 330.21 | 3.71 |
| *csp1* (ABUW_RS13055) | 325.75 | 15.48 |
| *ABUW_RS09460* | 322.14 | 3.07 |
| *raiA* | 318.52 | 3.65 |
| *ABUW_RS05005* | 293.8 | 60.17 |
| *tauA* | 242.59 | 16 |
| *ABUW_RS18750* | 211.35 | 3.98 |
| *tauB* (ABUW_RS11585) | 207.38 | 15.64 |
| *ABUW_RS09465* | 184.01 | 3.96 |
| *ABUW_RS04445* | 175.35 | 7.93 |
| *hscB* | 167.95 | 3.53 |
| *ABUW_RS16120* | 165.53 | 3.66 |
| *ABUW_RS17360* | 163.51 | 3.24 |
| *ABUW_RS20445* | 160.93 | 17.11 |
| *ABUW_RS10540* | 153.08 | 12.13 |
| *ABUW_RS01965* | 152.46 | 3.17 |
| *recX* (ABUW_RS08510) | 145.88 | 4.62 |
| *tauC* | 138.39 | 11.13 |
| *ABUW_RS18155* | 136.84 | 5.31 |
| *ABUW_RS07450* | 133.65 | 9.93 |
| *ABUW_RS09695* | 131.04 | 3.56 |
| *ABUW_RS13715* | 128.98 | 3.94 |
| *ABUW_RS18015* | 122.17 | 4.61 |
| *cysT* | 121.76 | 30.66 |
| *tauD* | 118.66 | 7.69 |
| *bfr_1* | 115.41 | 3.43 |

<sup>†</sup>Expression is reported as TPM of biofilm sample measured by RNA-seq<sup>‡</sup>Fold change is reported as expression of biofilm relative to planktonic sample

**Figure 2.**
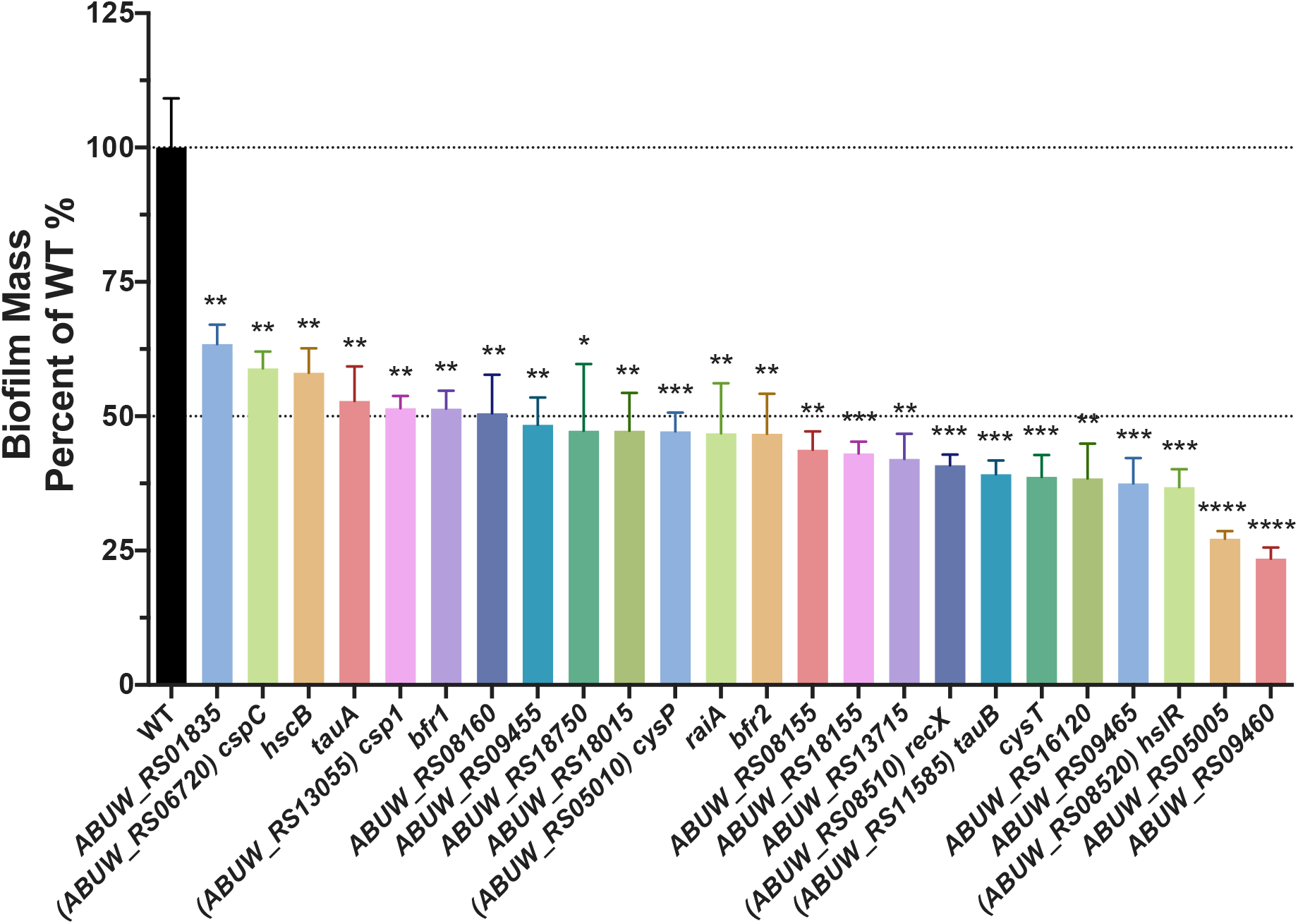
Physiological impact of gene disruption for those preferentially expressed in biofilms. Crystal violet biofilm assays were performed with transposon mutants of select genes preferentially expressed within biofilms. Alterations in biofilm mass are reported as a percentage of wildtype. Assays were performed in biological triplicate with 3 technical replicates each. Error bars represent ±SEM, Student’s *t* test was used to determine statistical significance. *, P≤0.05; **, P≤0.01; ***, P≤0.001; **** P≤0.0001.

To assess whether the observed changes in biofilm production may be due to a growth defect in the mutant strains, we measured cell density over time in liquid culture compared to wildtype (Fig. S2A). In so doing we noted that the vast majority of mutants behaved just as the wildtype, with only a handful of strains exhibiting growth defects. These included the transposon mutant for *csp1*, which grew in a similar manner to the parent for the first 3h, but then essentially plateaued in growth, reaching a maximum cell density (OD_600_) of 0.883±0.014 at 8.5 h (WT OD_600_ = 1.133±0.006 at 11.5 h). Another strain with a notable growth defect was the *tauA* mutant, which exhibited a less pronounced exponential growth phase, reaching a cell density of 0.602±0.011 by 4 h (WT OD_600_ = 0.769±0.018 by 4 h). The remaining mutant strains grew at rates comparable to wildtype and reached similar, if not higher, maximum cell densities. This indicates that the observed reduction in biofilm formation is unlikely to be an artifact of poor growth, with the exception of the *csp1* and *tauA* mutant strains.

### Disruption of *cspC* leads to changes in colony morphology and biofilm EPS

During our screen, we noted that one particular mutant, *cspC::tn* (+3.2-fold transcription in biofilm), produced a mucoidal, non-mucoviscous phenotype when grown on LB agar (Fig. 3). The *cspC* gene was one of the highest expressed genes within the biofilm (1324.94 TPM) and encodes a putative cold-shock protein, the role of which has not been previously characterized in *A. baumannii.* Importantly, complementation of *cspC in trans* restored colony morphology and biofilm production levels to those comparable to wildtype (Fig. 3 and 4A), and no measurable differences in growth (Fig. S2B). When observing *cspC* mutant cells by fluorescence microscopy, cell morphology was unaltered with respect to shape, size, and chaining (Fig. S3), indicating that obvious morphological differences are unlikely to contribute to the phenotype observed.

**Figure 3.**
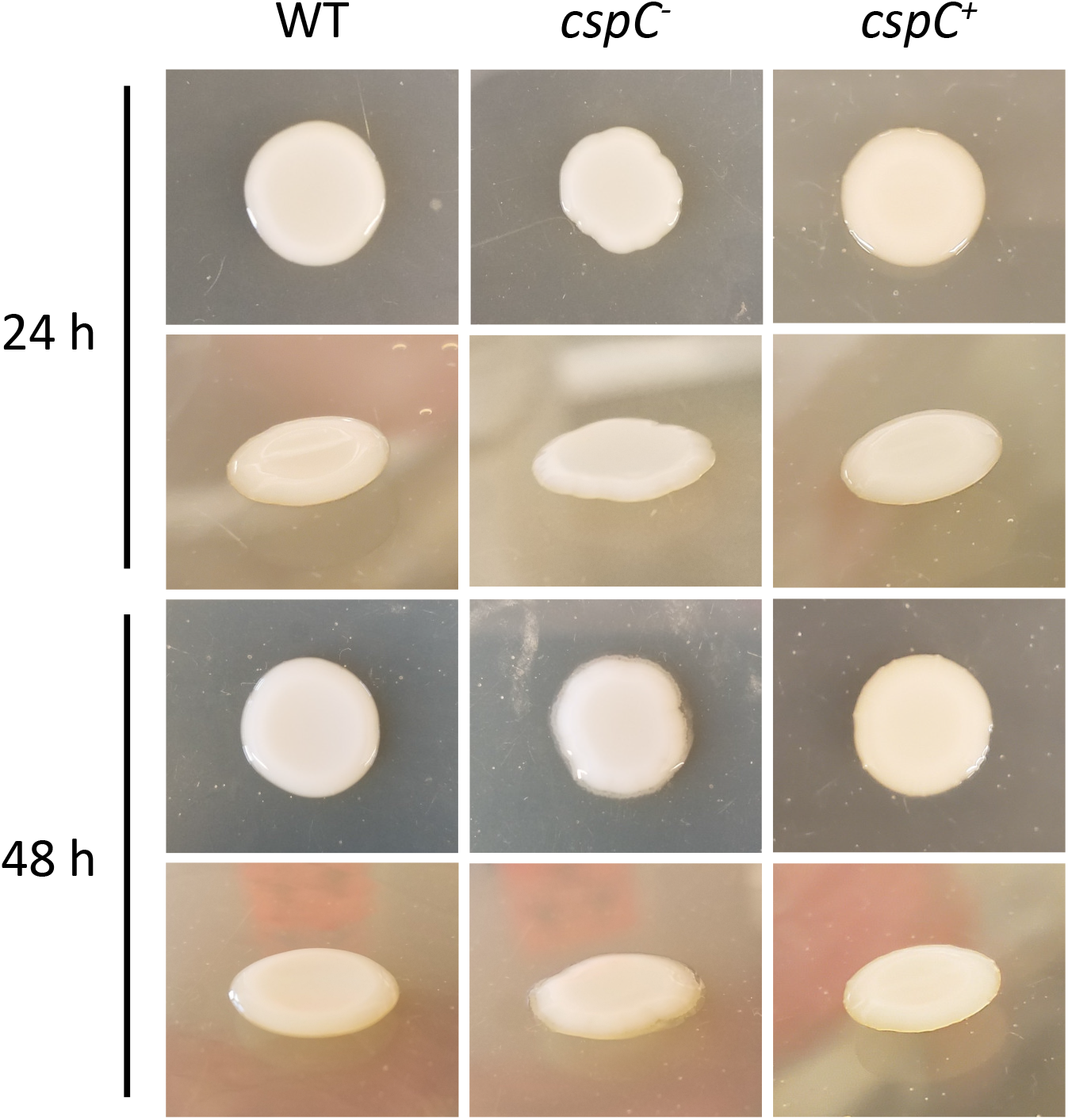
*cspC* mutants display an altered colony morphology. Wildtype (WT), *cspC^−^*, and *cspC^+^* strains were grown on LB agar supplemented with hygromycin at 37°C for 24 h (top) and subsequently left to grow for an additional 24h at room temperature (bottom). Images are representative of 3 experimental repeats.

**Figure 4.**
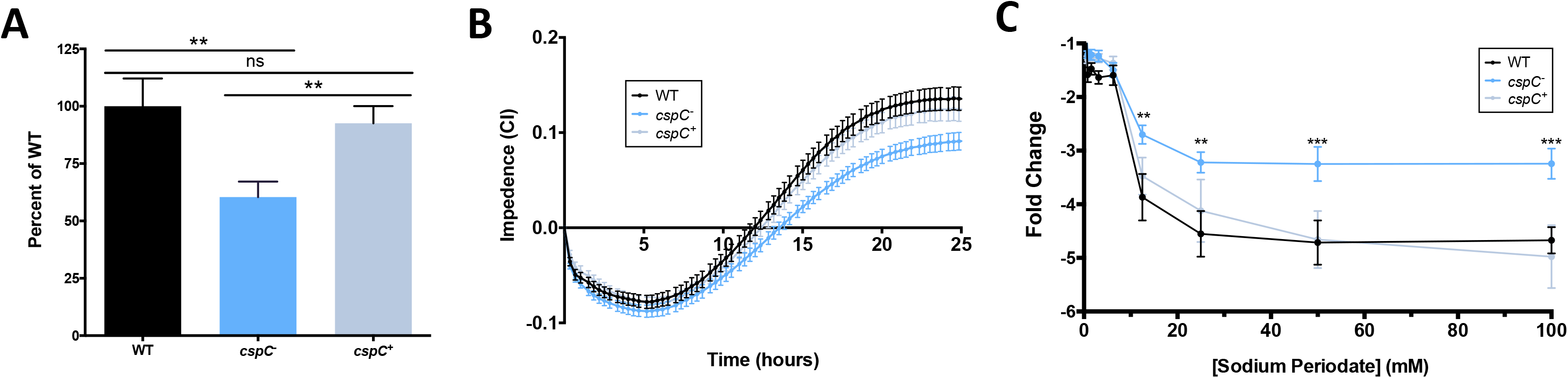
Disruption of *cspC* results in impaired biofilm formation and enhanced resistance to sodium periodate disruption. Comparison of biofilm formation using crystal violet staining following 24 hours growth (**A**), and continuously using an RTCA-xCELLigence (**B**) over a 25 h period. Assays were performed in biological triplicate with 3 technical replicates each. Error bars represent ±SEM; Student’s *t-*test was used to determine statistical significance compared to wildtype. **, P≤0.01, ns = not significant. For biofilm inhibition assay (**C**), biofilms were seeded with increasing concentrations of sodium periodate and incubated for 24h, with the resulting biofilms measured using a crystal violet assay. Data points are from 10 biological replicates and 3 technical replicates. Fold change is reported relative to no treatment controls for each strain. Error bars represent ±SEM. A two-way ANOVA was used to determine statistical significance from the wild type. **, P≤0.01; ***, P≤0.001.

To gain insight as to how and when CspC impacts biofilm formation, we used an xCELLigence RTCA to track biofilm formation in real time. The RTCA measures biofilm progression based on impedance of electrical signals between electrodes lining each well of a specialized 96-well plate, and expresses these measurements as a relative unit cell index (CI). CI measurements are influenced by cell adherence, secretion of <u>e</u>xtracellular <u>p</u>olymeric <u>s</u>ubstance (EPS), and cell spreading (70, 71). When CI values reach their maximum and remain steady, this signifies that the biofilm has entered maturation (70). In agreement with our crystal violet studies, the rate of increase and overall maximum CI value reached was noticeably lessened for the *cspC* mutant strain when compared to the parent and complement strains (Fig. 4B).

As cold-shock proteins have demonstrated importance in cell aggregation (31), extracellular polysaccharide production, and membrane fluidity (32), we hypothesized that the altered cellular morphology and decreased CI values of the *cspC* mutant may be due to changes in EPS production. Indeed, upon testing, we noted that *cspC* mutant biofilms showed increased tolerance to polysaccharide degradation by sodium *meta-*periodate compared to the wild type and complement strains (Fig. 4C). Conversely, the *cspC* mutant biofilm contained a comparable amount of eDNA to wildtype; and when we performed biofilm profiling assays in the presence of proteinase K, no change in biofilm integrity was observed (Fig. S4) suggesting that protein and eDNA components are not contributing to the biofilm defect observed upon *cspC* disruption.

### *cspC* is induced under cold-shock conditions but does not impact cold shock survival

CspC bares the same domain architecture as classic cold-shock proteins, which are defined as nucleic acid-binding proteins that are induced during temperature downshifts (34). To explore whether this is true of *cspC*, its transcription was measured during cold stress. Upon analysis we observed that *cspC* transcript levels were moderately increased compared to expression at 37°C (Fig. S5A) by approximately 1.5-fold. When the *cspC* mutant was challenged with cold stress for 1 hour, however, we did not observe any impact on cell viability (Fig. S5B). Furthermore, the *cspC* mutant also did not exhibit impaired survival during sustained cold stress (Fig. S5C). It is noteworthy that the *cspC* transcript is relatively abundant at 37°C, in both biofilm (1324.94 TPM) and planktonic conditions (408.71 TPM). Active transcription under varied conditions, coupled with the apparent lack of impact during cold stress, suggests that the role of CspC may extend beyond cold adaptation in *A. baumannii*.

### Transcriptional regulation by CspC

Csps are multifunctional proteins that can wield impacts at the level of transcription, post-transcription, and translation (reviewed in (34, 35)). How they exert their pleiotropic effects is either as a transcriptional regulator or RNA chaperone (33). To inform on the mechanism(s) governing the observed physiological changes upon *cspC* disruption in *A. baumannii*, we performed RNA-seq with our mutant and wild-type strains (Fig. 5A, Table S3). In so doing, we observed 147 genes upregulated and 54 genes downregulated by ≥2-fold upon *cspC* disruption (Fig. 5B). When sorted ontologically, it is apparent that a large portion of upregulated transcripts pertained to phage-related genes within the same chromosomal locus (n = 49) (Fig. 5C). Of these, 31 fall within the region predicted to encode Acinetobacter phage Bϕ-B1251, and the remaining 17 are within Ab105-1ϕ. Recent work has identified these regions as encoding active *Siphoviridae-* and *Myoviridae-*family phages, respectively, and as the two most prevalent phages identified among *A. baumannii* genomes (72).

**Figure 5.**
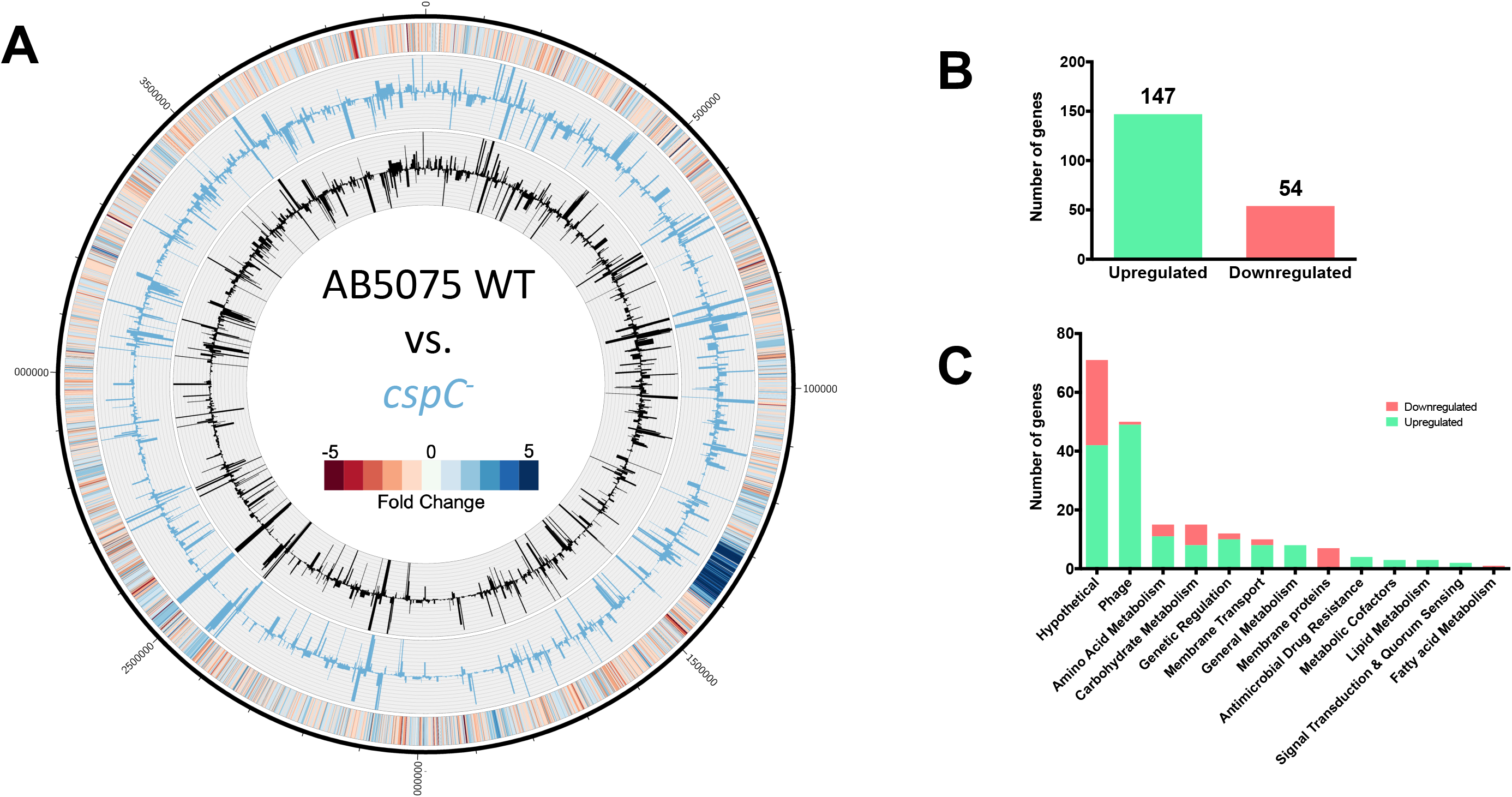
*A. baumannii cspC* disruption leads to global transcriptional changes. The genomic map (**A**) depicts transcription profiles of the wildtype and *cspC* mutant reported as TPM expression values. Inner histograms display RNA-seq expression values of the wildtype (black) and *cspC* mutant (blue) reported as TPM. The outermost circle is a heat map illustrating fold change in expression upon *cspC* disruption relative to wildtype, where blue or red indicates the fold-increase or -decrease of expression, respectively. The number of genes upregulated ≥2-fold (green) or downregulated (red) are tallied (**B**), and pa_5_rsed by function based on KEGG ontology (**C**).

A notable trend of opposing regulation of membrane transporter proteins compared to non-transporter membrane proteins was also apparent in our dataset. All 8 differentially expressed membrane protein genes, with no transport function, were downregulated, including type 1 pili subunits *csuAB* (6.19-fold) and *csuC* (2.7-fold), and fimbrial subunit, *fimA* (3.28-fold), each of which is important for attachment of *A. baumannii* to abiotic surfaces (24, 26, 73). Of the differentially expressed membrane transporters, 8 of 10 showed increased transcription, including *adeFG* (≥4.78-fold), which encode components of multi-drug efflux pumps. Also upregulated were the accompanying component *adeH* (3.29-fold) and additional efflux pump genes, *adeIJ* (≥2.08-fold) and *abeS (ABUW_RS06550*, 2.43-fold), which were categorized separately under Antimicrobial Drug Resistance by KEGG ontology. The AdeFGH efflux pump has been reported to confer resistance to DNA damaging agents and chloramphenicol, among other antimicrobials (74), and its overexpression has been correlated to increased biofilm formation of *A. baumannii* clinical isolates, by potentially serving a role in quorum-sensing (75). Like *adeFGH*, *adeIJK* and *abeS* encode components of multi-drug efflux pumps known to confer antimicrobial resistance in *A. baumannii* (74, 76, 77). Overexpression of *adeFGH* and *adeIJK* has previously been shown to decrease production of biofilm in *A. baumannii*, and for *adeIJK* overexpressing mutants, this has been attributed to decreased CsuAB, CsuC, and FimA abundance (78).

Other ontologies with enhanced expression in the *cspC* mutant included amino acid metabolism (e.g. – *pepD, glyA*); regulatory factors (e.g. – *rpoH, adeL, ABUW_RS06565*); and carbohydrate metabolism (e.g. – *gapN, atoA, atoD*). Collectively, it is clear that cspC has a global impact on the transcriptional homeostasis of *A. baumannii* and acts to regulate factors with many and varied roles within the cell.

To validate RNA-seq findings, a selection of genes were assayed by Real-Time Quantitative Reverse Transcription PCR (RT-qPCR). Trends in differential expression within the *cspC* mutant relative to wildtype were comparable to that of RNA-seq findings (Fig. S6).

### CspC influences the antimicrobial resistance profile of *A. baumannii*

Previous work employing a spontaneous mutation screen identified *cspC* as one of several genes able to restore, at least in part, antibiotic resistance in an AB5075 *bfmRS*-null strain (79). Considering this, and the upregulation of multi-drug efflux pump components observed in our study, we hypothesized that the *cspC* mutant would display phenotypic antibiotic tolerance. Indeed, previous work has shown that the AdeIJK efflux system (strongly upregulated in the *cspC* mutant) contributes to resistance towards a wide range of substrates including β-lactams, chloramphenicol, ciprofloxacin, and, in part, aminoglycosides (76, 80). Accordingly, we conducted an MIC screening panel for the *cspC* mutant using a variety of antimicrobial agents. In so doing we noted that disruption of *cspC* resulted in increased tolerance to ciprofloxacin, chloramphenicol, and streptomycin, yet slightly increased sensitivity to gentamicin, kanamycin, neomycin and fosfomycin (Table 3). Previous studies have demonstrated a strong correlation between *adeIJK* expression and ciprofloxacin resistance, with *adeIJK* overexpression mutants exhibiting enhanced resistance and *adeIJK* null strains demonstrating sensitivity; a phenomenon that is conserved across *A. baumannii* strains (74, 76, 78, 81, 82). Similarly, for multiple strains, transcription levels of *adeFGH* and *adeIJK* correlates with chloramphenicol resistance, with *adeFGH* transcription having the greatest influence (74, 76, 78). Thus, it seems possible that the upregulation of *adeIJK* in the *cspC* mutant is contributing to the measurable increase in ciprofloxacin tolerance, and similarly, the upregulation of both *adeIJK* and *adeFGH* may be contributing to enhanced chloramphenicol resistance. The disruption of *cspC* also resulted in slightly higher sensitivity to the majority of aminoglycosides tested (gentamicin, kanamycin, neomycin). Deletion of *adeABC* and *adeIJK* has been shown to induce gentamicin sensitivity, with *adeABC* deletion having significantly greater impact (78, 81, 82). In agreement with this, previous work has demonstrated that overexpression of *adeABC,* but not *adeFGH* nor *adeIJK*, enhances gentamicin, kanamycin, and neomycin resistance (78). Considering this, and that the *cspC* mutant has elevated *adeFGH* and *adeIJK* expression, but unchanged *adeABC* expression, the increased sensitivity to aminoglycosides is likely not attributed to these efflux pumps.

**Table 3.** Antibiotic susceptibility of AB5075 wildtype and *cspC* mutant strain (μg/mL).

| Strain | CIP | CM | STR | GM | KAN | NEO | FOS | OX |
| --- | --- | --- | --- | --- | --- | --- | --- | --- |
| AB5075 wildtype | 45 | 84.38 | 1070 | 2000 | 2250 | 253.13 | 562.5 | 1130 |
| AB5075 *cspC::tn* | 80 | 150 | 3380 | 1500 | 1690 | 189.84 | 316.41 | 1130 |

Abbreviations: CIP, ciprofloxacin; CM, chloramphenicol; STR, streptomycin; GM, gentamicin; KAN, kanamycin; NEO, neomycin; FOS, fosfomycin; OX, oxacillin.

### CspC regulates mRNA stability of transcriptional regulators *adeL* and *ABUW_RS06565*

When bacteria experience temperature downshifts, single-stranded DNA and RNA secondary structures are consequently stabilized. This leads to inhibited transcription and/or translation and RNA degradation, which adversely effects cellular function. Csps possess a conserved nucleic acid-binding domain that allows them to bind single-stranded DNA and RNA, and rescue undesirable secondary structures. Specifically, this nucleic acid-binding domain is typically comprised of a ribonucleoprotein (RNP)-1 and RNP-2 motif, which facilitates interaction with nucleic acids and has chaperone activity (34). When interrogating the amino acid sequence of CspC, a conserved cold shock domain signature was apparent (Fig. S7). This domain contains both a RNP-1 and RNP-2 motif (Fig. 6A). Structural prediction of CspC revealed that these motifs are both within the β2 and β3 strands of the antiparallel β-barrel structure (Fig. 6B). This arrangement makes the RNP motifs spatially available for binding to ssDNA and RNA, as is the hallmark characteristic of Csps (83, 84).

**Figure 6:**
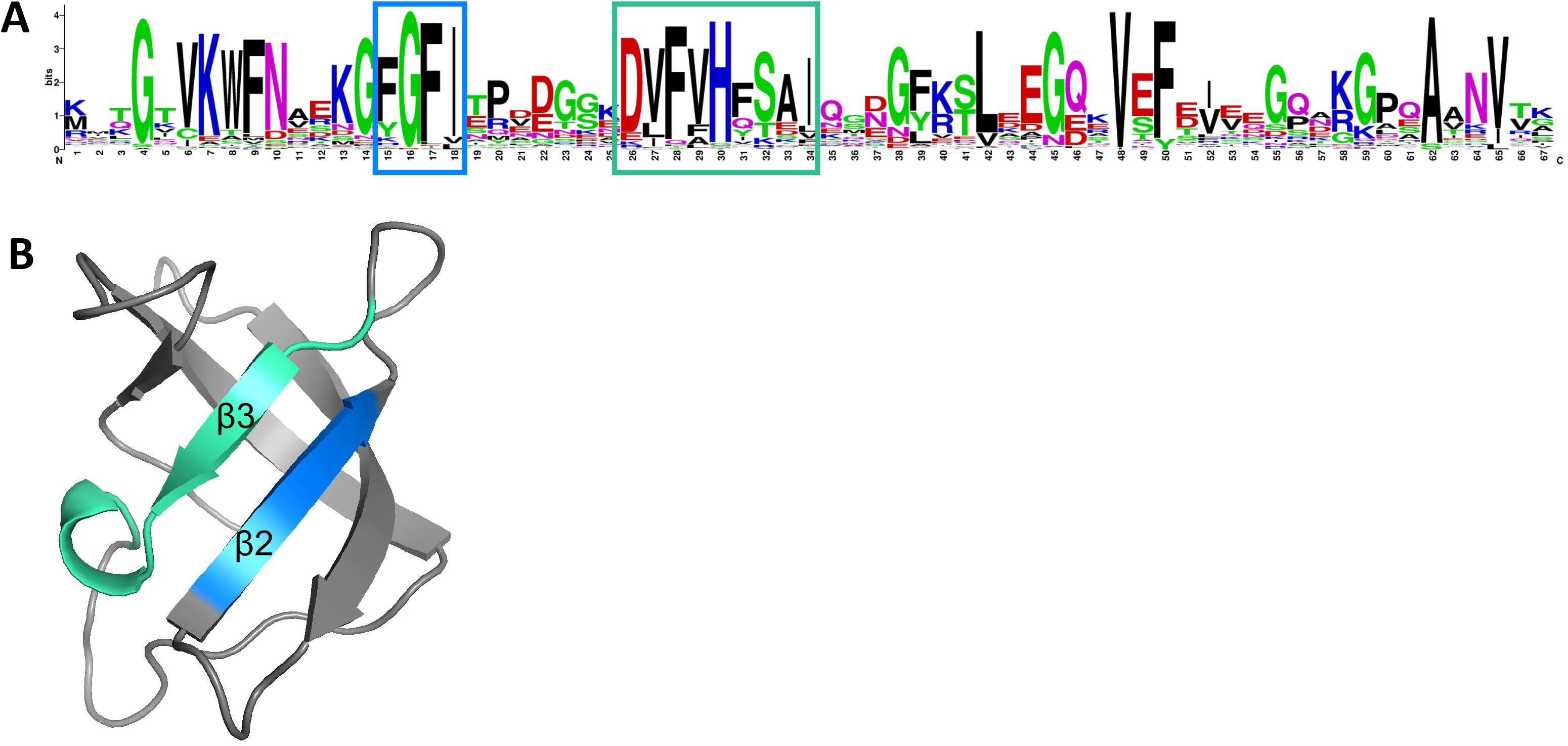
CspC contains conserved RNP-1 and RNP-2 motifs. A sequence logo (B) generated using the UniprotKB/Swiss-Prot sequences for the cold shock domain profile of AB5075 CspC detected by ScanProsite. Putative RNP-1 (blue) and RNP-2 (green) motifs are boxed. A 3-D representation of the *A. baumannii* CspC protein structure (B) is shown. Key residues within the β2 and β3 sheets are colored, corresponding with the RNP motif residues indicated in (A).

In order to determine if CspC may function as an RNA chaperone for the identified differentially expressed targets, we assessed the impact of *cspC* mutation on the rate of decay for putative target mRNAs. To do so, cells were grown to exponential phase and treated with rifampin to inhibit transcription. RNA was then isolated from wildtype and *cspC* mutant strains at consecutive time points post-transcriptional arrest. Using RT-qPCR, we assessed RNA transcript levels at each timepoint and determined decay rate by plotting the change in transcript abundance relative to transcript level immediately prior to rifampin treatment. The close arrangement of efflux pump encoding genes *adeFGH* indicates that these genes are likely within an operon, and similar conclusions can be drawn for *adeIJK*. Further supporting this, RNA-seq read alignments (data not shown) revealed transcript readthrough between the individual genes of each efflux pump. Considering this, we selected *adeG* and *adeJ,* located in the middle of their respective operons, for mRNA half-life studies. The half-life of *adeJ* mRNA was slightly longer than that of *adeG* (Fig. S8, Table 4) in the wild-type strain, however half-lives for both transcripts were comparable between the parents and *cspC* deficient strains. This suggests that CspC has no impact on the stability of *adeJ* and *adeG* mRNA. Importantly, the half-life of *recA* was also tested, and found to be 4.674 minutes in wildtype, which is comparable to the previously determined half-life of 4.5 minutes (57), indicating that the experimental approach used herein is reproducible.

**Table 4.** Mean mRNA half-lives of CspC-regulated transcripts as determined by RNA decay assays.

| Gene | Wildtype |  |  | *cspC::tn* |  |  |
| --- | --- | --- | --- | --- | --- | --- |
|  | Half-life (min)* | Decay constant (*k*)<sup>†</sup> | R<sup>2</sup> | Half-life (min)* | Decay constant (*k*) | R<sup>2</sup> |
| *adeJ* | 5.326 | 0.1301±0.015 | 0.954 | 5.206 | 0.1331±0.020 | 0.928 |
| *adeG* | 4.773 | 0.1452±0.016 | 0.926 | 4.236 | 0.1636±0.024 | 0.884 |
| *adeL* | 1.280 | 0.5417±0.020 | 0.999 | 2.540 | 0.2728±0.022 | 0.969 |
| *ABUW_RS06565* | 1.753 | 0.396±0.025 | 0.990 | 2.565 | 0.270±0.016 | 0.993 |
| *recA* | 4.674 | 0.1483±0.002 | 0.999 | 4.890 | 0.1418±0.031 | 0.880 |

\*The decay constant (*k*) mean was used to calculate half-life using $t_{1/2} = \ln(2)/k$ , where $t_{1/2}$ represents mRNA half-life.
<sup>†</sup>Decay constant is reported as mean ± standard error.

Considering that the half-lives of the major efflux pump transcripts were unaffected, we next considered whether CspC could be acting as an mRNA chaperone for transcriptional regulators. Transcription of *adeFGH* is repressed by the LysR type transcriptional regulator *adeL* (upregulated +2.8-fold in the *cspC* mutant, Table S3, Fig. S6), located immediately upstream of the *adeFGH* operon (74). When assessed experimentally, we found that the half-life of *adeL* was increased in our mutant from 1.28 minutes in the parent to 2.54 minutes in the *cspC^−^* strain (Fig. 7A, Table 4). Further to this, we also investigated the half-life of *ABUW_RS06565,* an uncharacterized GntR family transcriptional regulator, which was upregulated 2.98-fold in the *cspC* mutant. *ABUW_RS06565* was the only transcriptional regulator with altered transcription, aside from *adeL,* that was not part of the collectively altered Bϕ-B1251 and Ab105-1ϕ phage loci genes. The half-life of *ABUW_RS06565* rose from 1.75 minutes in the wild-type strain to 2.57 minutes in the mutant – a 1.5-fold increase in the *cspC* mutant compared to the parent (Fig. 7B, Table 4). This indicates that the mRNAs of transcriptional regulators *adeL* and *ABUW_RS06565* are in fact more stable as a result of *cspC* disruption.

**Figure 7.**
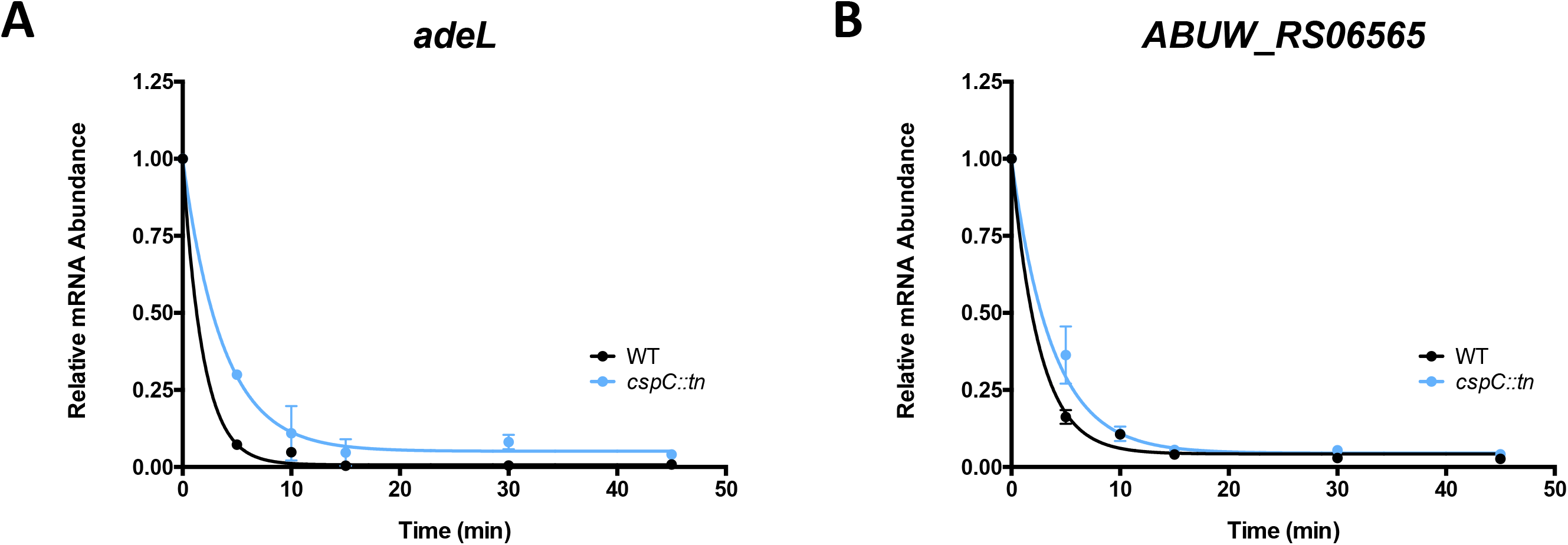
CspC mutation results in extended mRNA half life for transcriptional regulators, *adeL* and *ABUW_RS06565*. Exponentially growing *A. baumannii* wildtype and cspC mutant strains were treated with 250 µg/mL rifampin to arrest transcription and changes in transcript abundance was measured by RT-qPCR for *adeL* (A) and *ABUW_RS06565* (B). Values represent mean fold change in transcript abundance relative to transcript abundance immediately prior to rifampin treatment (t=0) ± standard deviation. Lines represent the exponential, one phase decay curve, represented as R(*t*) = R_0_*e^−kt^*, which were used to calculate mRNA half lives.

The average mRNA half-life varies depending on bacterial species and growth condition, however, the average RNA half-life hovers between 2-5 minutes (85–88). Considering this scale, the extended half-lives of *adeL* and *ABUW_RS06565* RNA in the *cspC* mutant are likely substantial. It is important to note that increased half-life of *adeL* does not necessarily indicate enhanced AdeL repression of *adeFGH*. Csps have the ability to destabilize RNA secondary structures in order to liberate the RBS and permit translation (reviewed in (35)), therefore, the absence of CspC could cause enhanced *adeL* stability resulting in increased transcription, but abrogated translation. In support of this notion, RNA secondary structure prediction revealed a stable hairpin formed in the 5’UTR of *adeL,* immediately upstream of a putative AGGAG ribosomal binding site (RBS) (Fig. 8). This is of particular interest as previous studies measuring mRNA half-lives in *E. coli* determined that an AGGAG motif in 5′UTRs, specifically within 2 to 8 nucleotides of translational start codons, is frequently found in transcripts with enhanced stability compared to the global mRNA population (89). The *ABUW_RS06565* transcript presents a curiosity as an obvious RBS is not apparent, however a similar, stable 5’UTR structure is also evident. Additionally, the *ABUW_RS06565* transcript contains an unusual, A/T-rich region beginning at the 5^th^ codon. A/T-rich sequences within RNA, particularly from the 5^th^ to 8^th^ codon of a protein coding region, have been shown to significantly enhance translation initiation in *E. coli.* This suggests that the inaccessibility of this motif due to enhanced RNA stability may also impede translation initiation of this transcription factor upon *cspC* deletion (90).

**Figure 8.**
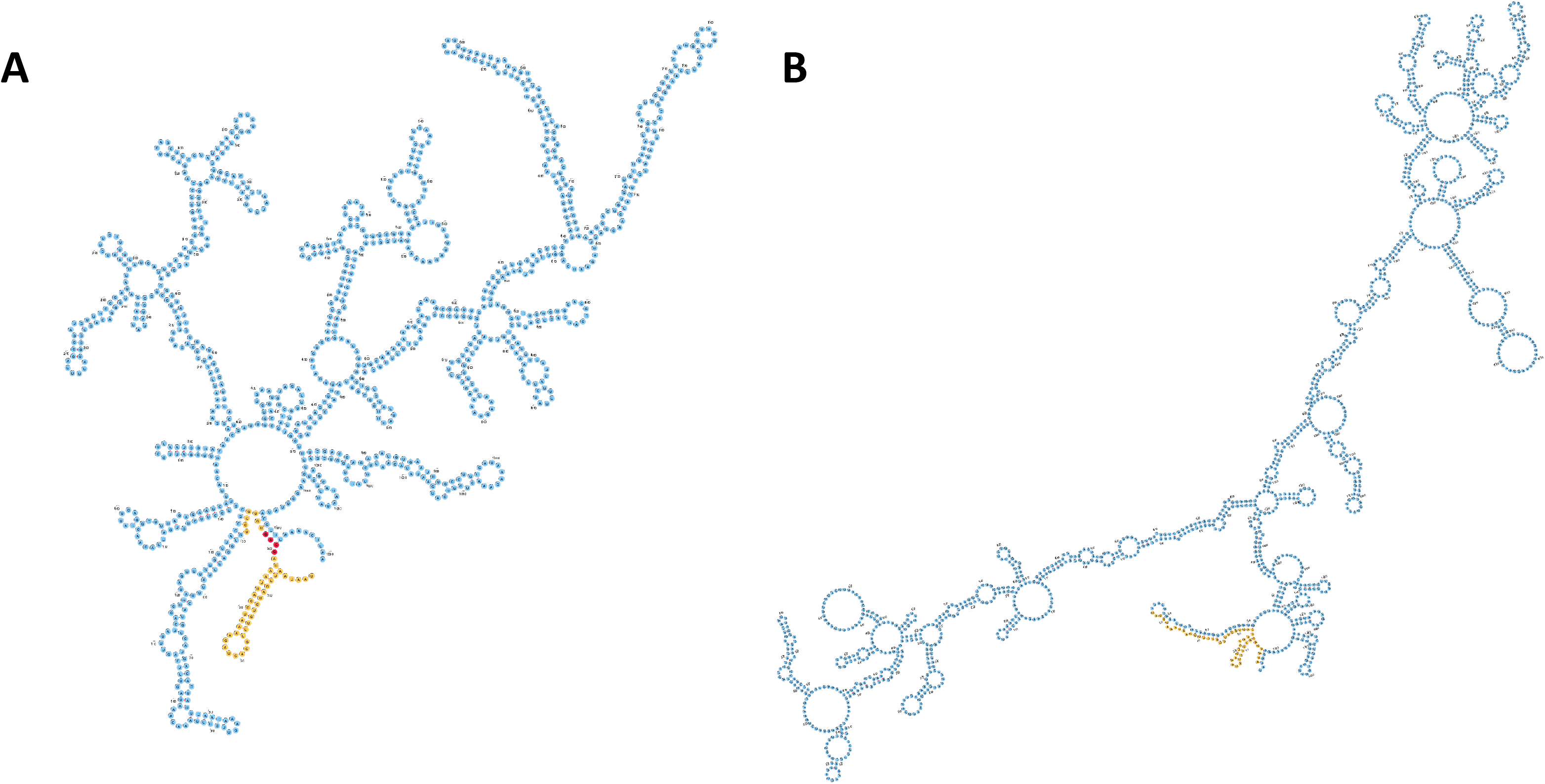
5’UTR of *adeL* mRNA contains a stable structure in close proximity to the putative ribosomal binding site. The secondary structure for *adeL* (A) and *ABUW_RS06565* (B) mRNA predicted using RNAfold is shown. Nucleotide colors correspond to the 5’UTR (yellow), putative ribosomal binding site (red), and protein coding sequence (blue). For *ABUW_RS06565,* a ribosomal binding site was not apparent.

### CspC is required for survival during oxidative stress and challenge by components of the human immune system

Biofilm production contributes to bacterial survival against host immune responses (91), and previous studies have found that a majority of *A. baumannii* bloodstream isolates are capable of producing robust biofilms (92). Furthermore, altering the balance of attachment, growth, and dispersal of a biofilm results in negative impacts on the capacity to cause bloodstream infections (93). Given that the *cspC* mutant has decreased propensity for biofilm formation, we speculated that the loss of CspC would impact *A. baumannii* survival in human blood. Thus, we grew the wild-type, mutant and complement strains in whole human blood for 6 h. Upon analysis we noted that AB5075 cell viability declined 67.6% from 0-1h, followed by an increase in viability as time progresses (Fig. 9A). In contrast, the *cspC* mutant strain fared markedly worse, demonstrating a far more severe decline in cell viability from 0-1h (86.6%), with cell viability counts remaining relatively stagnant as infections progressed. As expected, the *cspC* complement strain had a similar survival capacity as the wild type, indicating a clear requirement for CspC for survival during *A. baumannii* engagement with human blood.

**Figure 9.**
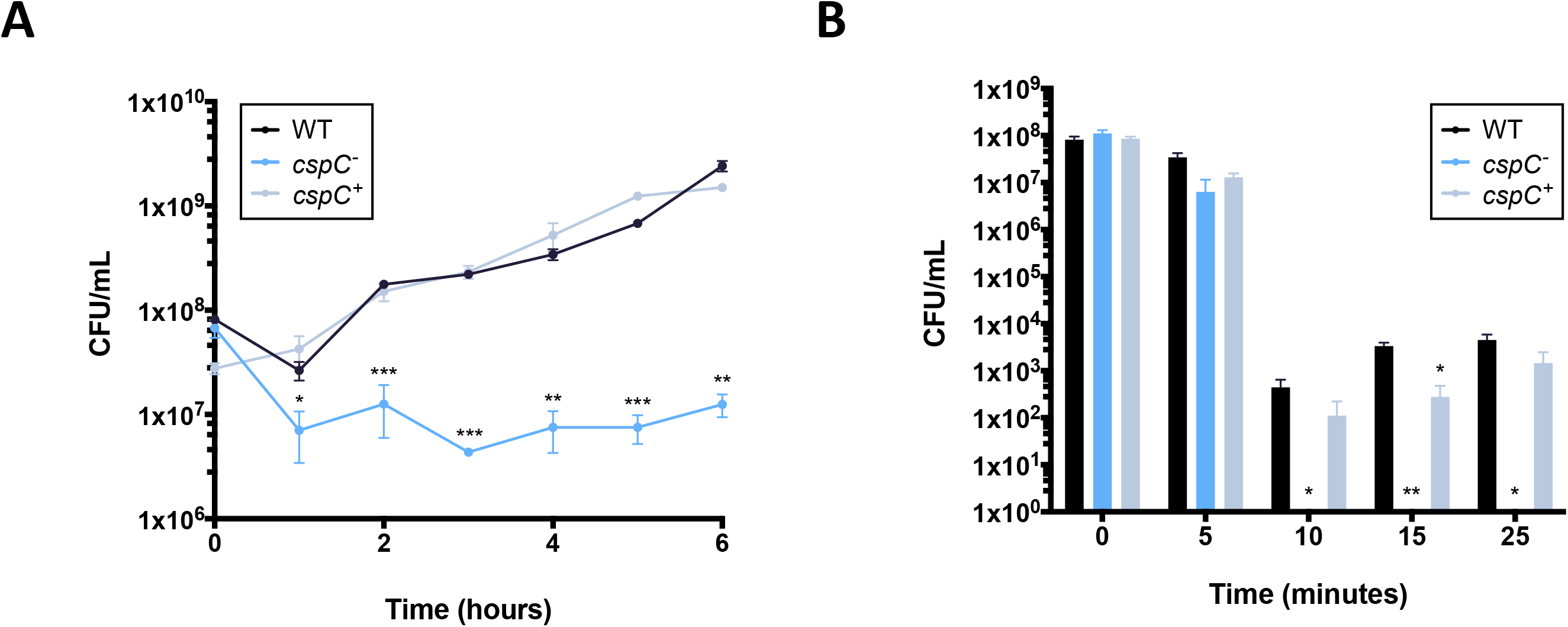
The *cspC* mutant has abrogated survival in human blood and during oxidative stress. Cell viability of the *cspC* mutant was assessed in whole human blood (**A**) and in the presence of 2mM hydrogen peroxide (**B**). Assays were performed in biological triplicate. Data is presented as the mean and error bars represent ±SEM. Student’s *t* test was used to determine statistical difference to wildtype at each timepoint. *, P≤0.05; **, P≤0.01; ***, P≤0.001.

Bacterial clearance in human blood is mediated by a number of factors, including cell mediated immunity. A primary killing mechanism of leukocytes during engagement with bacterial pathogens is wielded through reactive oxygen species (ROS). It has been previously demonstrated that clearance of *A. baumannii* is dependent on host reduction of nicotinamide adenine dinucleotide phosphate (NADPH) oxidase and subsequent accumulation of ROS (94, 95). Considering this, we hypothesized that the *cspC* mutant would likely have a survival defect upon exposure to oxidative stress, and thus performed a hydrogen peroxide killing assay (Fig. 9B). Upon analysis, we noted that the viability of all strains declined after 10 minutes of exposure to 2 mM hydrogen peroxide, which is a clear indication that the cells are experiencing oxidative stress. At conclusion of these studies (25 minutes) we observed a comparable number of viable cells were recoverable for the wildtype and complement strains. Conversely, the *cspC* mutant showed no cell recovery beyond 10 minutes of exposure, which indicates that CspC has a major role in bacterial survival for *A. baumannii* in the presence of ROS.

### CspC plays a pivotal role during *A. baumannii* systemic infections

Given our findings regarding the ability of the *cspC* mutant to survive challenge by components of the human immune system and reactive oxygen species, we next considered whether it’s survival during *in vivo* infection would be impaired. Accordingly, using a murine model of infection, we compared the pathogenic potential of the *cspC* mutant to wild-type AB5075. When assessing mortality, we observed no apparent differences between the strains (data not shown), however, when comparing dissemination of infection, bacterial load was substantially reduced for the *cspC* mutant strain in all organs evaluated (Fig. 10). The most significant reduction observed was for that of the liver, which showed a 43-fold reduction in bacterial load compared to wild type. Although not quite as striking, reduced dissemination of the *cspC* mutant to the other organs was equally remarkable (spleen: 12.16-fold; brain: 6.58-fold; lungs: 5.62-fold; kidneys: 4.32-fold, heart: 3.14-fold). These findings support CspC as a novel regulator of virulence, proving critical for *A. baumannii* survival when challenged by the host innate immune response.

**Figure 10.**
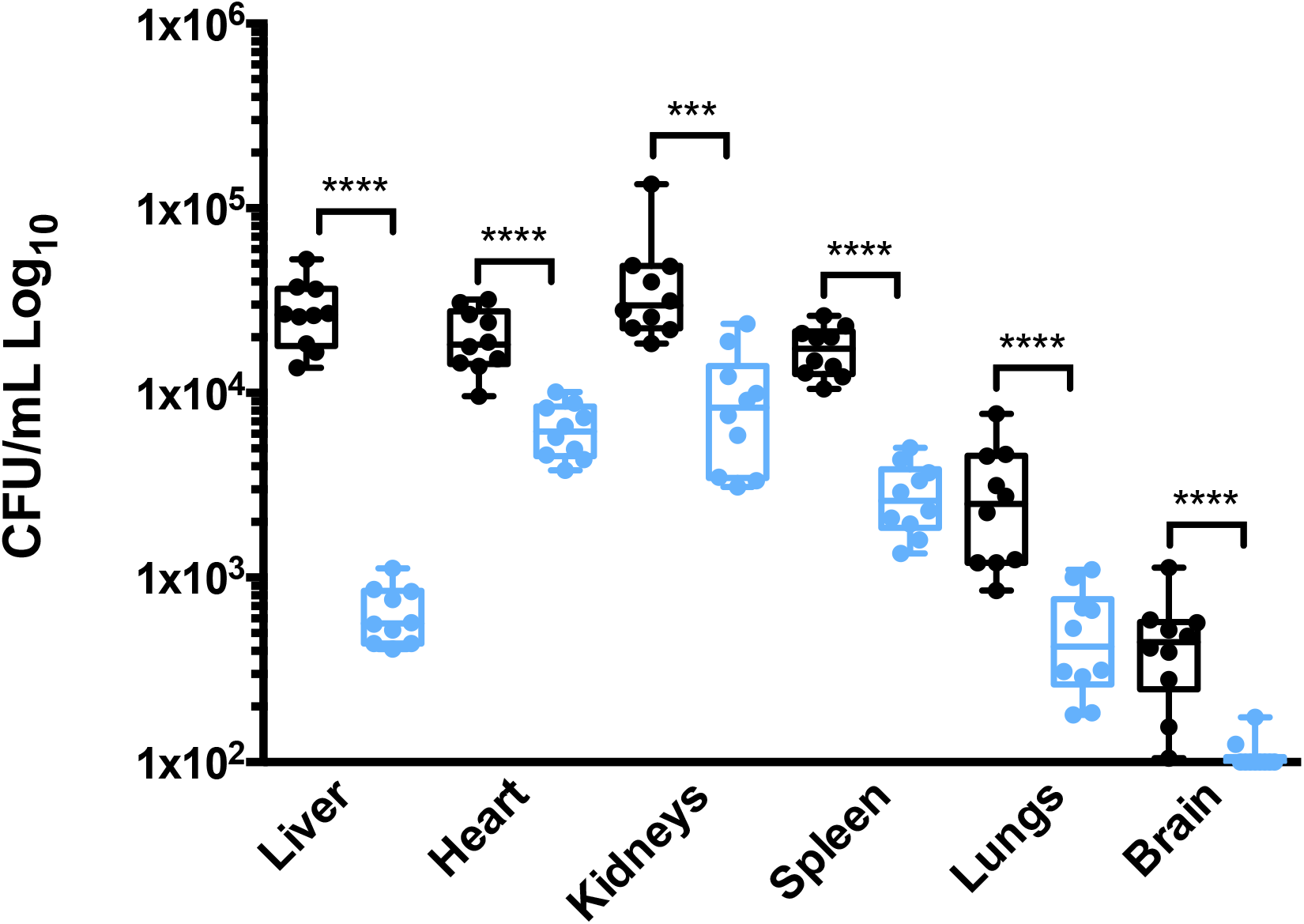
The *cspC* mutant is attenuated virulence for virulence in a murine model of sepsis. Cohorts of BL/6 mice (*n* = 10) were inoculated by retro orbital injection with 2.5×10^7^ CFU of wildtype (black) and *cspC::tn* mutant strains (blue). At 6 h post infection, mice were euthanized, and organs were collected to determine bacterial load. The bounds of each box represents 25^th^-75^th^ percentiles, whiskers represent 5^th^-95^th^ percentiles, center line denotes the median, and individual data points are plotted as circles. Statistical significance was determined using a Mann-Whitney nonparametric test. ***, P<0.001; ****, P <0.0001.

## Discussion

In this study we investigated the global transcriptional regulation occurring within biofilms formed by *A. baumannii* AB5075. RNA-sequencing revealed a distinct regulatory response within the biofilm, including the upregulation of 10% of all transcription factors identified to date, and the activation of sulfur metabolism pathway gene transcription. The most upregulated gene within biofilms compared to planktonic populations was *ABUW_RS05005* (+60.17-fold), which encodes an uncharacterized alpha/beta hydrolase fold protein co-expressed with the *cys* sulfate transporters (*cysP, cysT, cysW, cysA*). Flanking genes *cysP* and *cysT* were also significantly upregulated within the biofilm (+51.8-fold and +30.66-fold respectively). Microbiosis (0.1-0.3% O_2_) has been shown to induce sulfur metabolism gene transcription (96), and with anaerobic regions known to occur within biofilms (97, 98), conditions may be prompting a metabolic shift towards sulfur metabolism in our studies. *SsrS* was the most highly expressed gene within biofilms (62175.62 TPM), which produces the well-conserved, 6S RNA (69). Studies in *E. coli* have demonstrated that 6S RNA accumulates during stationary phase and represses transcription, in turn, enabling cell survival amid nutrient-limiting conditions (99, 100). It is quite possible that nutrient scarcity within the biofilm’s dense bacterial community may induce 6S RNA transcription to reorder gene expression circuits to circumvent nutrient limitation.

Among the enriched transcripts within the biofilm, we identified 24 which had significant physiological impact on biofilm integrity. Mutation of the most highly expressed of these (*cspC*) resulted in pleiotropic impacts on the cell. CspC is one of four known cold shock protein genes in AB5075, none of which have been thoroughly interrogated for physiological function. Interestingly, *cspC* was one of two Csps upregulated during biofilm growth – the other being *csp1* (+15.48-fold). The function of some Csps have been well studied under cold-shock conditions in a variety of organisms (101–103), however, Csps have diverse physiological impacts and Csp-mediated regulatory mechanisms, especially for those induced by non-cold shock conditions, remain poorly understood (reviewed in (33)). CspC transcription was induced under both cold stress and biofilm conditions; however, it did not have an appreciable role during cold shock. In biofilm, CspC seems to regulate extracellular polysaccharide production, as demonstrated by enhanced tolerance of the *cspC* mutant biofilm towards sodium *meta-*periodate. Extracellular protein and eDNA matrix components were unaffected by CspC, which is perhaps unsurprising as *A. baumannii* clinical isolate biofilms are primarily composed of polysaccharides (104). Increased tolerance to polysaccharide degradation may indicate overproduction and/or structural variation of this matrix component. If the former were true, we would anticipate enhanced biofilm production, but this was not the case. We suggest that structural variation, which would render polysaccharides resistant to sodium *meta-*periodate, may be impairing biofilm integrity in the *cspC* mutant. Such a scenario is supported by the observations of others, where altered colony morphology and abrogated biofilm formation as a consequence of polysaccharide modification has been well described for a variety of bacteria (reviewed in (19)).

In previous work, spontaneous mutation of *cspC* bypassed antibiotic hypersensitivity and cell morphology defects in a *bfmRS* mutant strain, which was attributed to reduced transcription of the well-known capsule factor - the K locus (79). No transcriptional change for *cspC* was evident in the *bfmS, bfmR,* or *bfmRS* mutant strains (79), and in our work, *bfmRS* transcription was unaffected by *cspC* disruption. Given that mutation of *cspC* is able to bypass sensitivity phenotypes in previous work, and that the *cspC* mutant has enhanced tolerance to polysaccharide degradation, we anticipated upregulation of the K-locus, a major determinant of capsular polysaccharide production in *A. baumannii*. Surprisingly, no measurable changes for any genes within the K locus were observed, indicating CspC may act independently from the capsule biosynthesis factors within the K locus. Instead, our findings suggest that CspC may mediate biofilm formation through regulation of the *csu* pili assembly system and *fimA* fimbrial subunit transcription, both of which have demonstrated roles in the attachment of *A. baumannii* to abiotic surfaces and were downregulated in the *cspC* mutant strain (24, 26, 73). In particular, Csu pili are adhesive organelles that belong to the archaic chaperone-usher pili class, which facilitate strong adherence to hydrophobic plastics, including polypropylene and polyethylene, which are widely used in medical equipment (26) and are indistinguishable from the materials used in this study. The *cspC* mutant also exhibited elevated multi-drug efflux pump component (*adeFGH, adeIJK)* transcription. Overexpression of *adeIJK* has previously been shown to decrease production of CsuAB, CsuC, and FimA, in turn, altering membrane composition and decreasing biofilm formation in *A. baumannii* (78). In addition to biofilm abrogation, we suspect the cooperative activation of *adeFGH* and *adeIJK* may be contributing to enhanced chloramphenicol resistance as their contribution to resisting the effects of this drug has been well documented (74, 76, 78). Similarly, heightened *adeIJK* expression in the *cspC* mutant may be responsible for the marked increased in ciprofloxacin resistance, as studies have demonstrated a strong correlation between *adeIJK* expression and resistance to this particular fluoroquinolone (74, 76, 78, 81, 82). The consequential upregulation of efflux pumps upon *cspC* deletion may explain why antibiotic resistance was restored for the antibiotic-sensitive *bfmRS* mutant strain once *cspC* mutation was introduced in previous work (79).

Interestingly, transcription of *adeFGH* repressor, *adeL* (74), was also upregulated in the *cspC* mutant. Csps can have impacts at the transcriptional, post-transcriptional, and translational levels (reviewed in (34, 35)) and we suspect CspC is facilitating *adeL* translation and degradation. In support of this model, the putative *adeL* RBS is in close proximity to a stable secondary structure that likely occludes translation initiation; and *adeL* decay was slowed in the *cspC* mutant. CspC may thus bind the *adeL* mRNA, consequently destabilizing its secondary structure, liberating the RBS, and permitting RNA turnover. Under this model, CspC disruption would lead to increased *adeL* mRNA, but reduced AdeL abundance. Such a regulatory mechanism may not be limited to *adeL*, as the uncharacterized transcriptional regulator, *ABUW_RS06565*, also showed increased transcript abundance and a reduced rate of mRNA decay in the *cspC* mutant. *ABUW_RS06565* does not have an obvious RBS, however, its 5’UTR contains a similar stable secondary structure encompassing an A/T rich, translation-enhancing motif (90). It is quite possible that CspC serves a similar role in destabilizing *ABUW_RS06565* to facilitate translation.

Structural investigation revealed that CspC contains a conserved RNP-1 and RNP-2 nucleic acid binding motif, a hallmark characteristic of Csps. Interestingly, the RNP-2 motif contains an arginine residue at position 34, where most bacteria contain a serine residue. Furthermore, the RNP-1 and RNP-2 motifs are bridged by six amino acids, as opposed to the usual seven. The shortened motif separation, but not arginine residue substitution, is a common occurrence for all four *A. baumannii* cold shock proteins (CspC, ABUW_RS12225, Csp1, ABUW_RS15360), with ABUW_ RS15360 having the shortest bridge of 4 amino acids. The combination of both abridged RNP motifs and residue substitution in RNP-2, may alter the affinity of CspC to nucleic acid ligands.

Importantly, biofilm production contributes to bacterial survival against host immune responses and previous studies have found that a majority of *A. baumannii* bloodstream isolates produce strong biofilms (91, 92). Furthermore, altering the balance between attachment, growth, and dispersal of a biofilm reduces the ability to cause bloodstream infections (93). Disruption of *cspC* resulted in impaired survival in human blood likely due to its limited biofilm forming capacity, and inability to endure in the presence of ROS, the manifestation of which is one of the earliest *in vivo* defense mechanisms against *A. baumannii* (67, 68). Likewise, attenuated dissemination of infection was evident for the *cspC* mutant in a murine model, which establishes CspC as a critical *A. baumannii* virulence factor.

In summary, this study identifies CspC as having a critical role for biofilm formation, antibiotic resistance, and virulence via regulation of adhesins and multidrug efflux pumps. Additional studies are needed in order to determine if CspC acts as an RNA chaperone, as we have proposed, to modulate translation. Furthermore, this study identified 23 additional genes with substantial influence over *A. baumannii* biofilm, many of which remain uncharacterized and have not been previously implicated with biofilm formation. Future work will seek to understand how these genes contribute to the complex regulatory network governing *A. baumannii* biofilm formation and to assess their potential as therapeutic targets.

### Ethics statement

All animal work was performed under the approval of the University of South Florida’s Institutional Animal Care and Use Committee (IACUC).

## Acknowledgements

This study was supported by grants AI124458 and AI157506 (LNS) from the National Institute of Allergy and Infectious Diseases and GM133617 (PJE) from the National Institute of General Medical Sciences. The funders had no role in study design, data collection and interpretation, or the decision to submit the work for publication. We extend our thanks to the USF Genomics Program Genomics Equipment Core for the use of their facilities for RNA sequencing.

## Author contributions

Conceptualization: B.R.T., L.N.S.; investigation: B.R.T., G.D., R.S.B., P.J.E.; methodology: B.R.T, J.L.A; formal analysis: B.R.T.; writing – original draft preparation: B.R.T.; writing – review and editing: B.R.T., L.N.S.; Funding: L.N.S.

## Conflicts of interest

The authors declare that there are no conflicts of interest.

